# The genome of a subterrestrial nematode reveals an evolutionary strategy for adaptation to heat

**DOI:** 10.1101/747584

**Authors:** Deborah J. Weinstein, Sarah E. Allen, Maggie C. Y. Lau, Mariana Erasmus, Kathryn C. Asalone, Kathryn Walters-Conte, Gintaras Deikus, Robert Sebra, Gaetan Borgonie, Esta van Heerden, Tullis C. Onstott, John R. Bracht

## Abstract

The nematode *Halicephalobus mephisto* was originally discovered inhabiting a deep terrestrial aquifer 1.3 km underground. *H. mephisto* can thrive under conditions of abiotic stress including heat and minimal oxygen, where it feeds on a community of both chemolithotrophic and heterotrophic prokaryotes in an unusual ecosystem isolated from the surface biosphere. Here we report the comprehensive genome and transcriptome of this organism, identifying a signature of adaptation: an expanded repertoire of 70 kilodalton heat-shock proteins (Hsp70) and avrRpt2 induced gene 1 (AIG1) proteins. We find that positive selection has driven the expansion of Hsp70 genes, which are also transcriptionally induced upon growth under heat stress. We further show that AIG1 may have been acquired by horizontal gene transfer (HGT) from a rhizobial fungus. Over one-third of the genes of *H. mephisto* are novel, highlighting the divergence of this nematode from other sequenced organisms. This work sheds light on the genomic strategies of adaptation to heat in the first complete subterrestrial eukaryotic genome.

## Introduction

*Halicephalobus mephisto* was discovered inhabiting a fluid-filled aquifer accessed from the Beatrix Gold Mine in South Africa at 1.3km below the surface^1^. Radiocarbon dating indicates the aquifer water is over 6,000 years old^1^, and the lack of surface ^3^H infiltration, a remnant of atmospheric atomic testing, highlights its isolation from the surface biosphere^1^. The water is warm (37°C), alkaline (pH 7.9), hypoxic (0.42 - 2.3mg/L dissolved O_2,_), and rich in biogenic methane (CH_4_)^1–3^. In spite of these challenging conditions, a thriving, complex microbial community exists in this extreme environment including chemolithoautotrophic organisms that extract energy from the subterrestrial rock and fix organic carbon^2,4^. Syntrophic relationships link sulphur-oxidizing denitrifying bacteria, sulfate reducers, methanogens and anaerobic methane oxidizing organisms into a complex mutually reinforcing microbial food web^2^ that supports a rich assemblage of eukaryotic opportunistic predators including nematodes, rotifers, and protists^5^. With the exception of *H. mephisto* none of these eukaryotic organisms have been cultured in the laboratory, and none have had their genomes sequenced and analyzed until now.

Nematodes encode small, remarkably dynamic genomes well suited to studies of adaptation^6,7^. Among the most abundant animals on earth, nematodes have adapted to an incredibly diverse set of environments: from hot springs to polar ice, soil, fresh and saltwater^8^, acid seeps^9^, and the deep terrestrial subsurface^1^, with a wealth of comparative genomic data available. Dynamic gene family expansion^6,10^ and shrinkage^11^ have proven good signatures of evolutionary adaptive selection in crown eukaryotes, including nematoda^12^. Here we have performed comprehensive genomic and transcriptomic studies in *H. mephisto*, giving a first view of the evolutionary adaptive response to a subterrestrial environment, and have identified expanded gene families and patterns of expression under heat stress.

## Results

*De novo* DNA sequence assembly with Illumina data and scaffolding with PacBio reads, yielded a small complete assembly of 61.4 Mb comprising 880 scaffolds with N50 of 313kb, though 90% of the sequence is encoded on just 193 scaffolds. The longest scaffold is just under 2.55 Mb (Table 2). Several lines of evidence suggest that this is a highly complete genome. The Core Eukaryotic Genes Mapping Approach (CEGMA)^13,14^ identified 240 of 248 core eukaryotic genes for a completeness of 97%, and tRNscan-SE^15^ identified 352 tRNAs encoding all 20 amino acids plus selenocysteine. Benchmarking Universal Single Copy Orthologs (BUSCO)^16^ estimated the completeness as 81.4%, but manual inspection of its output shows that 79 “missing” genes are actually detected, with good e-values (median 6e-19). Given that BUSCO’s thresholds are established from 8 nematode genes from Clades I, III, and V,^17^ it may not be well suited to divergent Clade IV nematodes such as *H. mephisto,* since it also scored the *P. redivivus* genome (98% complete^18^) as only 82.1% complete. Therefore considering these divergent matches to be valid, we conclude that for *H. mephisto* BUSCO detected 946 / 982 orthologous genes, for a completeness of 96%, consistent with CEGMA. Reinforcing the completeness of the *H. mephisto* assembly and annotation, a quantitative comparison of 3,252 protein domains shows a strong correlation with *C. elegans* (Figure 1B).

**Figure 1.**
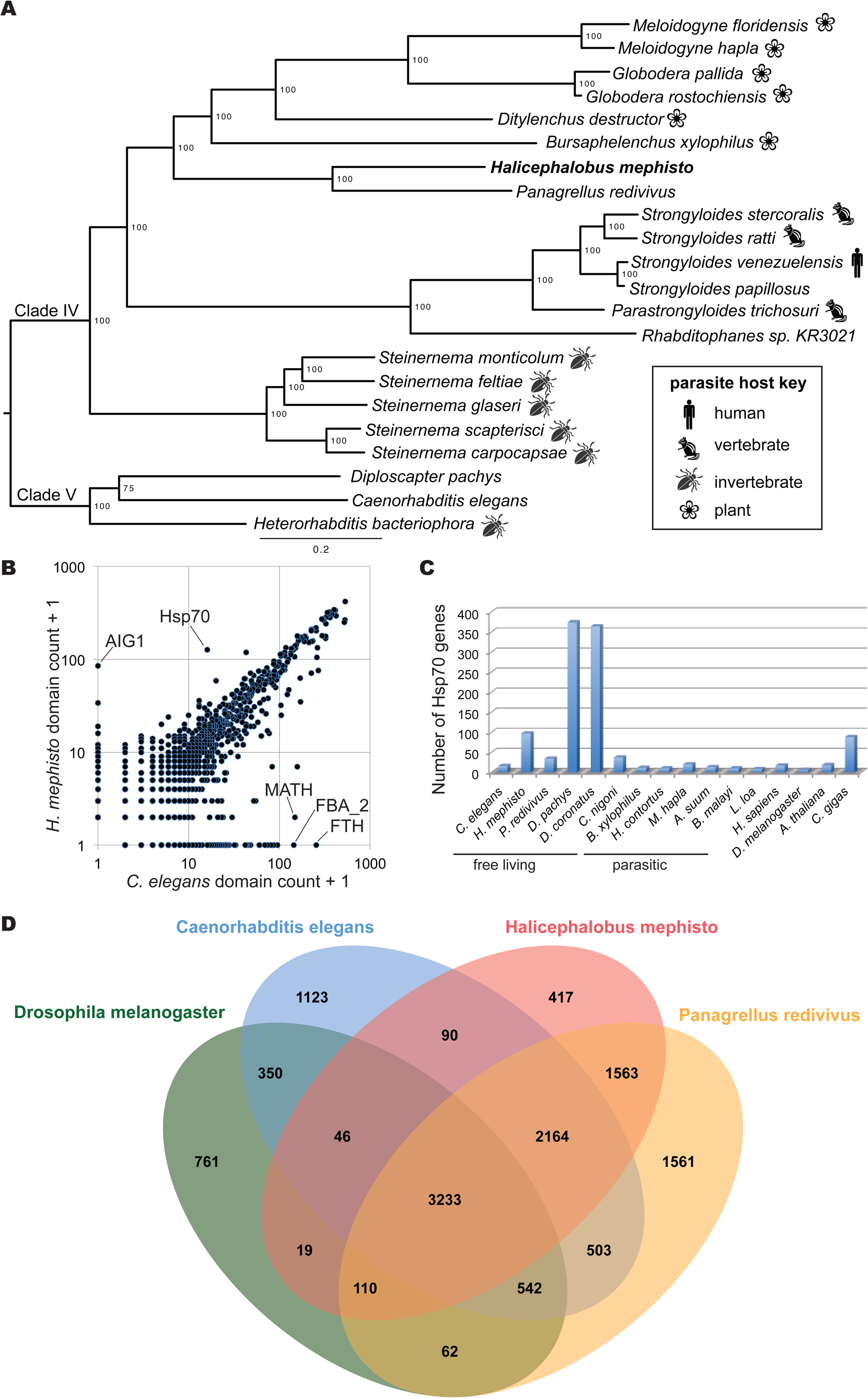
Genomic comparison of *H. mephisto* protein-coding genes. A. Multi-loci phylogeny of *H. mephisto* using 99 single-copy orthologous genes (SCOGS). B. Pfam domain comparison of nonredunant protein domain content in *C. elegans* versus *H. mephisto* at an e-value cutoff of 1e-10. Expanded AIG1 and Hsp70 domain families are marked, along with the MATH, FTH, and FBA_2 domains known *C. elegans*-specific expansions. C. Examination of Hsp70 family expansion across species, quantifying proteins, not domains, identified using e-values as in B. D. Venn Diagram comparing orthologous gene clusters between *D. melanogaster, C. elegans, P. redivivus* and *H. mephisto*. Abbreviations: MATH: Meprin And TRAF-Homology domain, FTH: Fog-2 Homology domain, FBA_2: F-box associated domain.

The repetitive component of the *H. mephisto* genome is highly novel. Using a custom RepeatModeler repeat library, RepeatMasker masked 24.3% of the genome, denoting 21.1% as interspersed repeats, 87.3% of which are unknown (Table 1). Evaluation of these sequences with nhmmer^19^ and the DFAM database positively identified 44.3% of these as helitrons, with 8.8% retrotransposons and 23.6% DNA transposons (Table 1). However, consistent with the genomic divergence of *H. mephisto,* this repetitive element repertoire appears extremely different from known elements, including many unique or novel repeat families needing further characterization, and a significant 23.3% remain unclassified by either algorithm (Table 1).

**Table 1.** Repetitive element composition of the *H. mephisto* genome. The percentage of repeats belonging to each class are shown.

|  | RepeatMasker | nhmmer |
| --- | --- | --- |
| Retrotransposons | 3.8 | 8.8 |
| DNA Transposons | 8.8 | 23.6 |
| Helitrons | 0.0 | 44.3 |
| Unclassified | 87.3 | 23.3 |

The *H. mephisto* nuclear genome was annotated with Maker2^20^, TopHat^21^, StringTie^22^, and TransDecoder^23^, and transcripts were clustered with gffcompare to define a total of 16,186 protein-coding loci. An additional 1,023 loci were identified that do not encode proteins over 50 amino acids and are candidate noncoding transcripts, for a total of n=17,209 transcribed loci in the *H. mephisto* genome, producing 34,605 transcripts for an average of 2.0 transcripts per locus (Table 2). Intron lengths averaged 473 compared to 320 bp for *C. elegans* (Table 2).

**Table 2.**
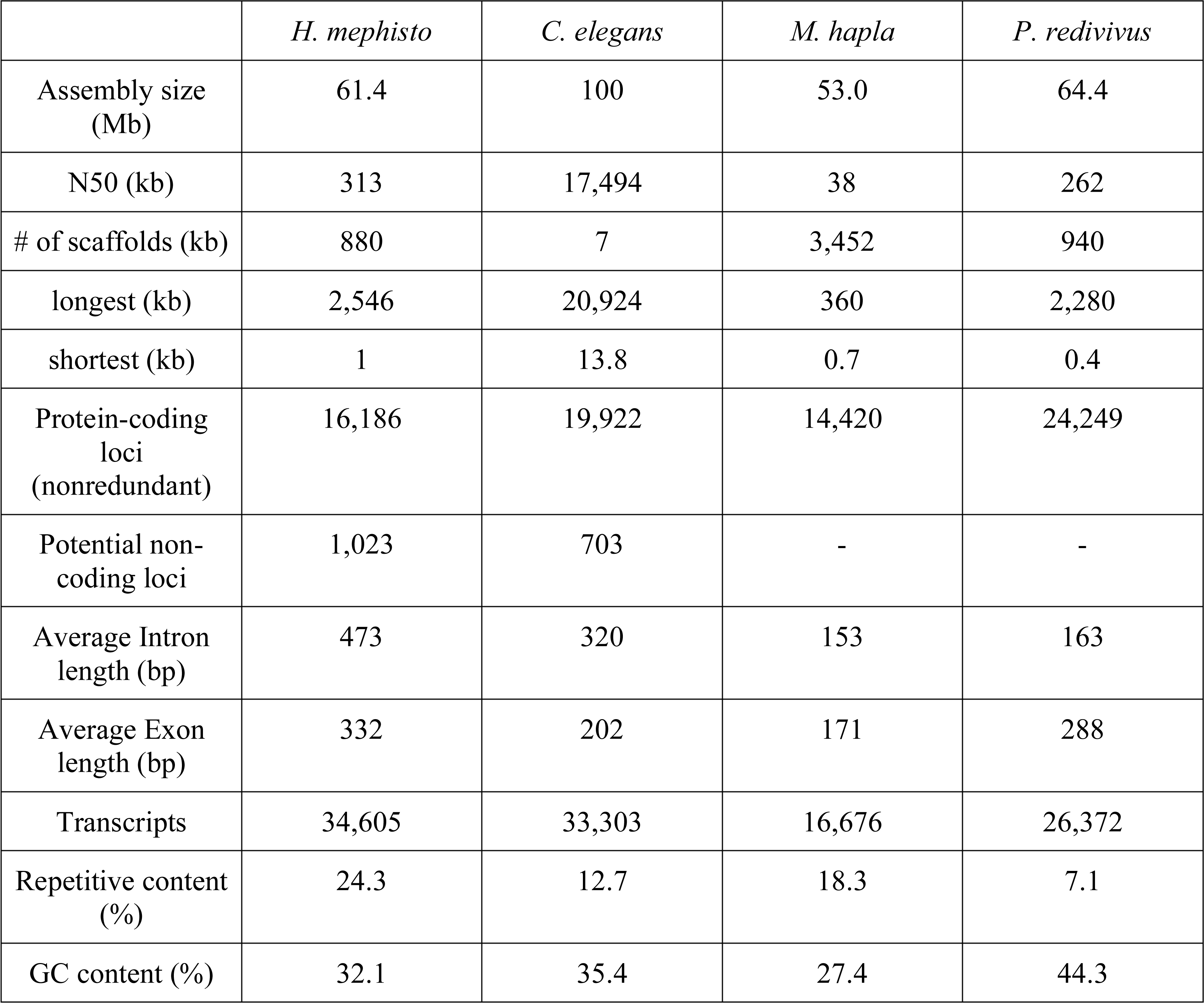
Genome and transcriptome data for *H. mephisto* and comparison to *C. elegans*, *M. hapla*, and *P. redivivus*. Data for *C. elegans* and *P. redivivus* from Srinivasan *et al*. 2013 and for *M. hapla* from Foth *et al*., 2014.

We used single-copy orthologous proteins (SCOGS) to build a phylogenetic tree placing *H. mephisto* as a distant relative of free-dwelling *Panagrellus redivivus*, the nearest fully sequenced nematode relative, within Clade IV (Figure 1A). The comparison of domain counts between *C. elegans* and *H. mephisto* uncovered two domains strikingly enriched in *H. mephisto*: *<u>a</u>vrRpt2*-induced <u>g</u>ene, AIG1 (84 domains vs. 0) and 70-kilodalton heat-shock protein, Hsp70 (126 domains vs. 15) (Figure 1B). The most over-represented domains in *C. elegans* are the Meprin And TRAF-Homology (MATH) domain, Fog-2 Homology Domain (FTH) and F-box associated (FBA_2) domains, all apparent lineage-specific expansions in *Caenorhabditis*^24,25^ (Figure 1B).

We used OrthoVenn^26^ to identify orthologous genes between *D. melanogaster*, *C. elegans*, *P. redivivus*, and the 16,186 *H. mephisto* proteins, identifying a set of 5,397 shared among all nematodes, with 3,233 shared among all four invertebrates. Among the 417 *H. mephisto*-specific orthologous groups, the largest cluster was Hsp70, with 107 proteins. The expansion of Hsp70 is particularly evocative because these well-studied heat-activated chaperones re-fold proteins denatured by heat^27–30^ as part of a coordinated response to heat-shock^31–34^. Even more intriguing, Hsp70 are expanded in organisms adapted to environmental thermal stress, but not in nematodes parasitic on endothermic hosts, suggesting the expansion of Hsp70 may be a general strategy for adaptation to environmental, not parasitic, heat (Figure 1C). Bayesian phylogenetic analysis of Hsp70 proteins recovered known paralogs specific to cellular compartments including mitochondrial and endoplasmic reticulum (ER)^35^ along with a cluster grouping human, mouse, and nematode genes (Cluster I, Figure 2A). The recovered Hsp70 gene tree topology is robust, given that the same structure was recovered by maximum likelihood (Figure S1). The human sequences in Cluster I include well-characterized Hsp70 sequences^36^. Cluster II is a new 37-member *H. mephisto-*only group, which surprisingly is most closely related to another novel *Diploscapter* cluster with 59 genes (Cluster III, Figure 2A). These data suggest that the Hsp70 gene family has undergone significant amplification within the *Diploscapter* and *Halicephalobus* lineages which, owing to their evolutionary distance (Figure 1A), most likely did not inherit these expanded gene families from a common ancestor. Instead, we propose these genes underwent independent expansions in both lineages under shared evolutionary pressure to adaptat to heat stress: *D. pachys* is thermotolerant^37^, while *D. coronatus* is a facultative parasite of humans^38^ and a member of the genus has been found in thermal waters^39^.

**Figure 2.**
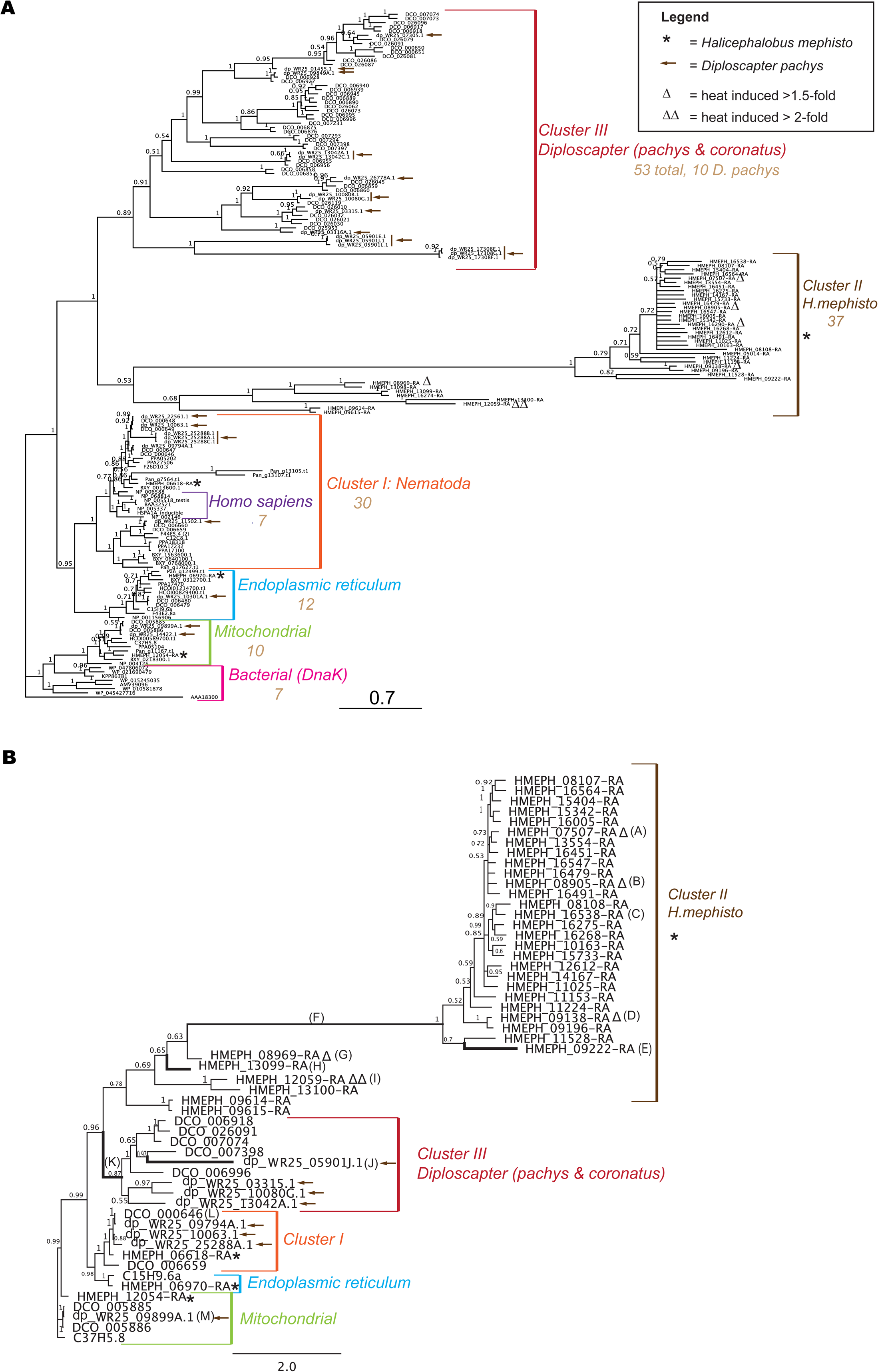
Analysis of Hsp70. (A) Bayesian phylogenetic tree of Hsp70. *H. mephisto* sequences marked with an asterisk (*) and *D. pachys* with arrows. Branch numbers indicate posterior probabilities, scale bar represents substitutions per site. (B) Hsp70 protein Bayesian tree used for dN/dS analysis of coding sequence. Branch letters A-M correspond to Table 2 for ω (dN/dS) value. Bold branches indicate statistical significance of dN/dS p-value after correcting for multiple hypothesis testing, and the long branch is semi-bold because it is statistically significant prior to multiple testing correction with 89% of sites under positive selection (Table 2). Branch numbers indicate posterior probabilities, scale bar represents substitutions per site. The sequences used in both trees are indicated with Wormbase (nematode) or Genbank Accessions (other species).

A signature of adaptive evolutionary change is positive selection, in which mutations altering amino acids (dN) are enriched relative to (presumably neutral) synonymous changes (dS), giving a dN/dS ratio (ω) greater than 1^40^. A value of ω less than 1 indicates elimination of mutations that alter the amino acid sequence, also known as purifying selection, a hallmark of evolution working to preserve protein function^40^. Given the long branch lengths and potentially selection-driven expansions of Hsp70 genes, we used PAML^41^ to test for positive selection on this gene family. To facilitate this analysis we created a new Bayesian phylogeny using a subset of genes from *H. mephisto*, the *Diploscapter* species and *C. elegans* outgroups, resulting in a well-resolved Bayesian phylogeny (Figure 2B). We ran a branch-sites test in PAML, which estimates two ω parameters for different codons on a pre-selected “foreground” branch of the phylogenetic tree. The ω_1_ parameter accounts for pervasive purifying selection at specific sites (codons) but a second ω_2_ measures values greater or equal to 1 (from neutral to positive selection) at other sites. Because ω_2_ is estimated from the data, it is an indicator of positive selection if it is reported to be above 1. Furthermore, by performing test twice, once with ω_2_ fixed at 1 (neutrality, a null model) and once allowing it to be freely estimated from the data, a likelihood ratio test can be used to derive a p-value quantifying the strength of positive selection on a particular branch^42^. In our analysis, positive selection was detected along the long branch leading to the *H. mephisto* cluster: ω_2_ of 197 on 89% of amino acids, p = 0.04 (Table 3), however it was not robust to Bonferroni correction for multiple hypothesis testing, which may be too conservative^43^. Given that this branch has by far the most sites (89%) under positive selection, we have made it semi-bold (Figure 2B). Values of ω_2_ greater than 1 were detected on ten of eleven branches from clusters II and III (Table 3) with four giving a p-value that is significant after correction for multiple hypothesis testing (lines strongly bolded in Figure 2B). In particular, the root of the *Diploscapter* cluster (Cluster III) showed evidence of strong positive selection (Table 3, Figure 2B). As a control, we tested a Cluster I short branch (L) and mitochondrial gene (M), which showed no evidence of positive selection (Figure 2B, Table 3). Together these data suggest that positive selection has driven the diversification of Hsp70 genes in both *Halicephalobus* and the *Diploscapter* lineages.

**Table 3.**
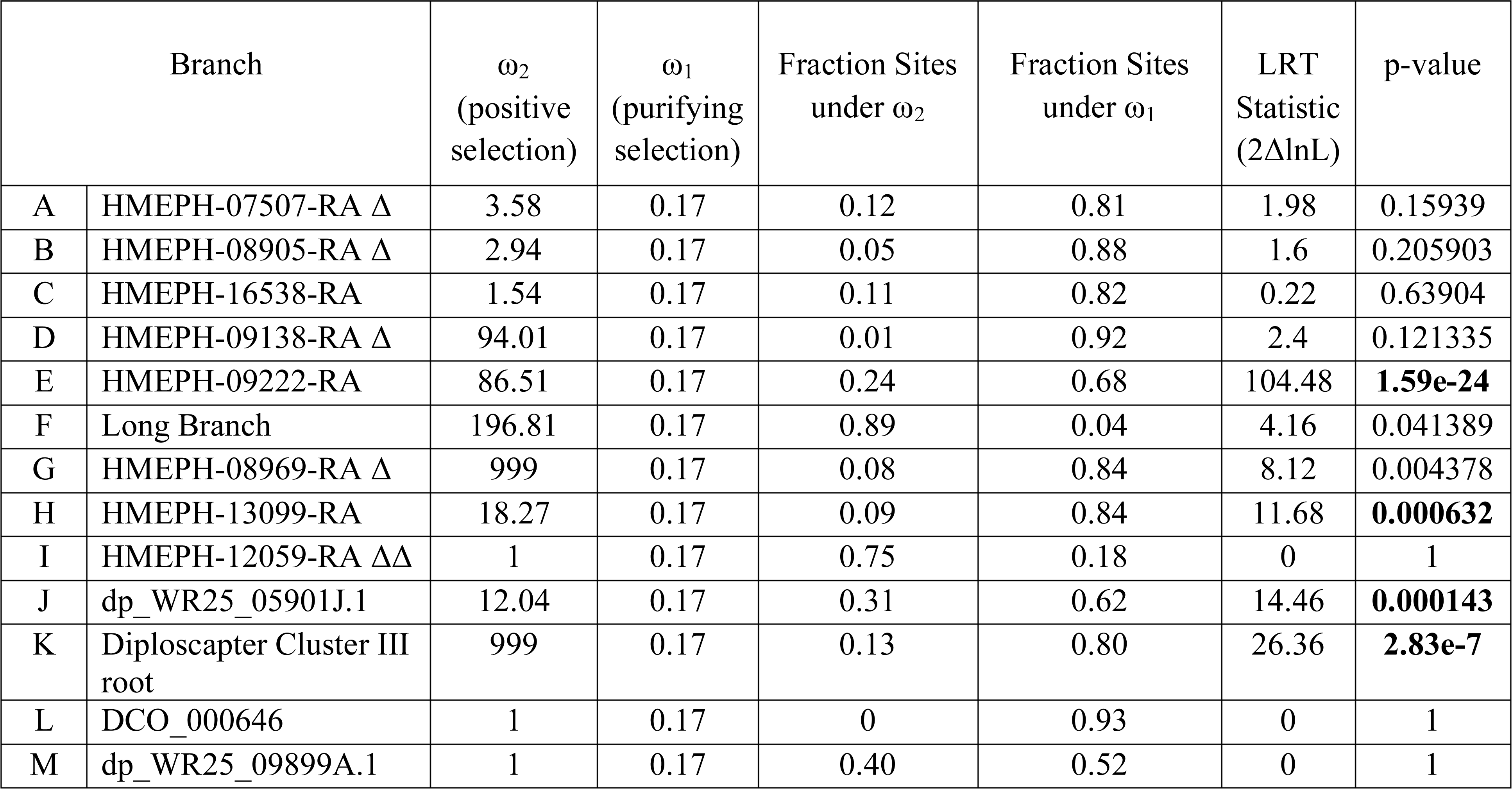
Branch-site analysis of dN/dS ratios (ω) from tree in Figure 2B. Bold p-values are statistically significant after correcting for multiple hypothesis testing (p=0.0038). Likelihood ratio test (LRT) statistic applied to a χ^2^ table with df=1, critical values 3.84 (5%) and 6.63 (1%). Note that ω_2_ is constrained to be greater than or equal to 1 (positive to neutral selection). Δ, expression increased 1.5x under heat stress. ΔΔ, expression increased 2x under heat stress.

AIG1 was originally identified as a pathogen response gene in plants^44^, but is also involved in survival of T-cells in mammals, where they were named immune-associated nucleotide binding proteins, (IANs)^45^ or GTPase of immunity-associated proteins (GIMAPs)^46^. These proteins function as GTP-binding molecular switches controlling cell fates^46^. AIG1 was originally reported to be completely absent from nematodes^45^, but by relaxing the statistical stringency we find a single copy of the domain (Y67D2.4) in *C. elegans* annotated as a homolog of human mitochondrial ribosome associated GTPase 1, MTG1 suggesting a possible divergence and expansion of the GTPase superfamily in *H. mephisto* and other nematodes. Consistent with this, blastp against the nr database and HMMER search of Uniprot reference proteomes identified hundreds of matches with low percent identity (∼30%) to the AIG1s identified in *H. mephisto.* Nonetheless, the matches range from 250-300 amino acids in length and with e-values from 1e-20 to 1e-30; they come from species as diverse as nematodes, fungi, arthropods, and the parasitic protist *Giardia intestinalis*. These sequences are generally annotated as uncharacterized or hypothetical proteins, though some are annotated as p-loop containing nucleoside triphosphate hydrolases, suggesting a large GTPase protein family, previously uncharacterized, resides within eukaryotes.

Consistent with the relatively low percent identity of these genes, most did not align well with the *H. mephisto* sequences, and if they did align, they did not show good bootstrap support in phylogenetic trees (data not shown). This suggests the eukaryotic superfamily of GTPases is comprised of distinct and divergent subfamilies, only one of which is the AIG1 domain expanded in *H. mephisto*. We ultimately were able to obtain good alignments and well-supported phylogenetic trees from only 17 nematode species: an AIG1-like cluster including *H. mephisto, Diploscapter coronatus, D. pachys, A. suum,* and *P. redivivus*, while a separate MTG1-related cluster includes the *Caenorhabditis* sequence and other nematodes (Figure 3A). Notably, the two distinct clusters resolve with 100% bootstrap support, and while the MTG1-related cluster includes members of all sequenced nematode clades (I,III,IV, and V; no Clade II genomes are currently available^47^), the AIG1-like group includes only members of clades III, IV, and V, suggesting a potential origin in the Chromadoria^47^ and extensive amplification in *H. mephisto* (Figure 3A).

**Figure 3.**
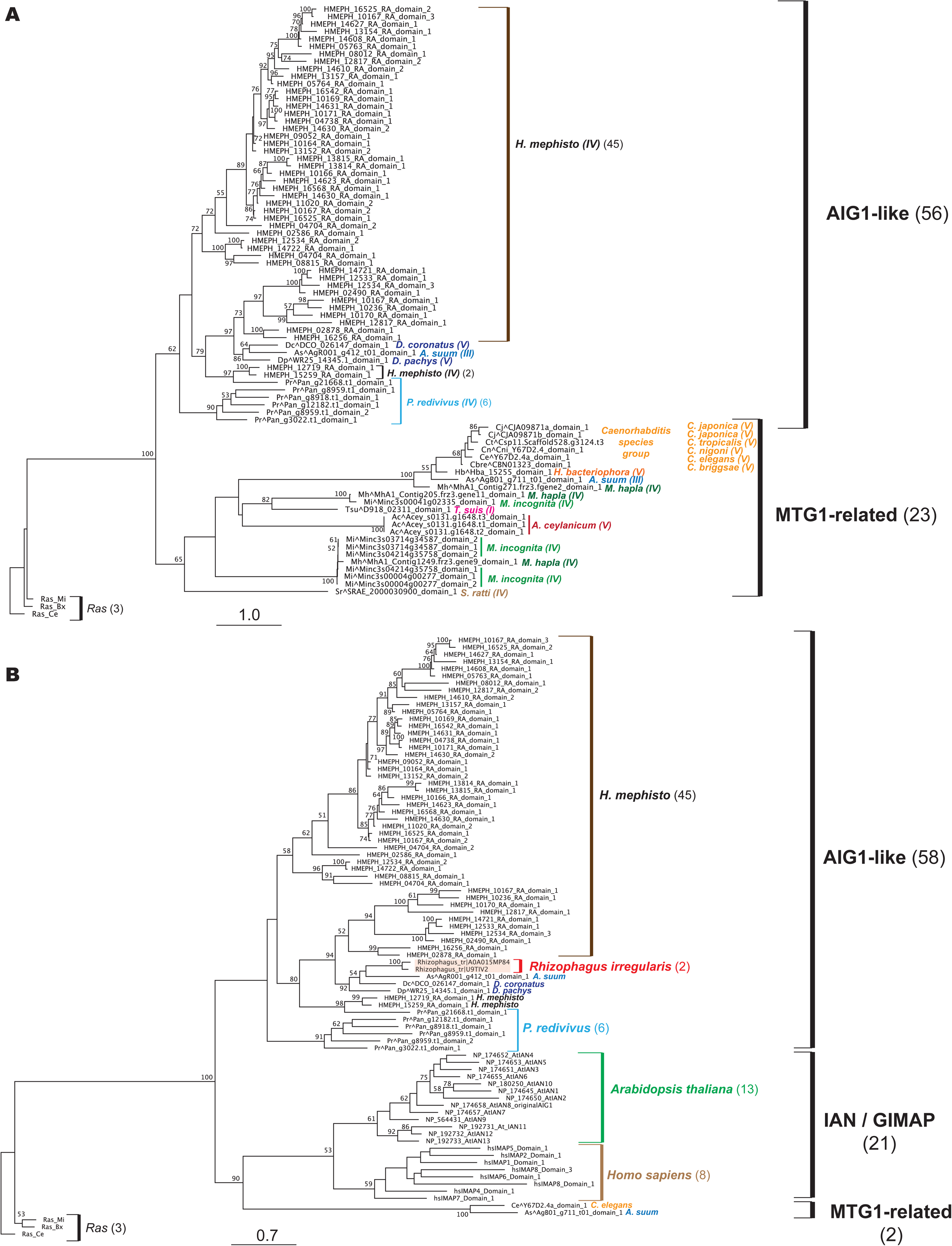
Phylgenetic analysis of AIG1. (A) Nematode-only RAxML tree of 17 species showing two clusters: MTG1-related and AIG1-like groups. (B) Nematode, human, *Arabidopsis*, and fungal RAxML tree illustrating potential HGT event. IAN / GIMAP are synonyms of AIG1, the original names given to plant and vertebrate sequences^46^. Two fungal (*Rhizophagus irregularis*) AIG1-like sequences highlighted in light red boxes. For both trees, branch numbers indicate bootstrap support from 200 replicates, and scale bar represents substitutions per site. The sequences used in both trees are indicated with Wormbase (nematode), UniProtKB (*Rhizophagus irregularis*) or Genbank Accessions (other species).

In order to relate these nematode GTPase subfamilies with the previously identified AIG1 / IAN / GIMAP sequences from vertebrates and plants^45,46^ we built a tree including a broader taxonomic sampling beyond nematodes (Figure 3B). To simplify the tree we included only six nematodes in this analysis, five which we previously characterized as having the AIG1-like genes: *H. mephisto*, *P. redivivus*, *A. suum, D. coronatus,* and *D. pachys,* and the *C. elegans* representative of the MTG1-related proteins. In this tree the original IAN/GIMAP human-plant cluster was recovered^45,48^ but deeply rooted from the MTG1-related side of the phylogeny with good branch support (Figure 3B). We found two sequences from *Rhizophagus irregularis*, a plant-root associated fungus, resolve cleanly into the newly discovered AIG1-like group including *H. mephisto* (Figure 3B). The *R. irregularis* sequences are from two single-nucleus genome sequences, accessions EXX68593.1 and EXX68414^49^. These data suggest that, in contrast to reports of AIG1 being absent in invertebrates^45,50^ Clade III, IV, and V nematodes do have a previously unknown member of this protein family potentially derived from an ancient horizontal gene transfer from a rhizobial fungus. *D. pachys* has been reported to inhabit the rhizobial zone of plant roots^37,51^, where it is ideally situated to acquire horizontally transferred genes from cohabiting fungus. Given that we found five nematode species host orthologs of the AIG1-like domain in their genomes (Figure 3A) the HGT event must have been in a Chromadorean ancestor lineage, not in contemporary *D. pachys*, though *H. mephisto* has undergone a spectacular expansion of this gene family that is not present in the other lineages, most of which have only a few copies (Figure 3A). The divergence of suborders Rhabditina (*C. elegans, D. pachys, D. coronatus*) and Tylenchina (*P. redivivus* and *H. mephisto*) has been estimated at 22 million years ago using molecular clock methods^52^. *A. suum* is a member of Spirurina and separated from the Rhabditina 80 million years ago^53^, and the putative HGT event must predate these divergences. While inter-domain HGT into eukaryotes has been controversial, transfer from fungus to *C. elegans* has been documented, setting a precedent for this hypothesized HGT event^54^. Preliminary analysis of codon bias to test for HGT was performed, but caution has been urged given this method’s propensity for misleading results^55^; indeed while we found the codon usage of AIG1 genes differed statistically from the rest of *H. mephisto* coding sequences, this was also true for our control datasets including collagens, tubulins, and even Hsp70 (data not shown).

We performed a comprehensive differential gene expression analysis of *H. mephisto* genes whose expression changes under heat stress, identifying 285 heat-upregulated and 675 heat-downregulated transcripts (Figure 4B). Because many upregulated genes were unknown, and the upregulated set is small, Gene Ontology analysis identified no enriched categories. However, the downregulated set of genes was enriched in peptidases, as well as cuticle components and ornithine-oxo-acid transaminase (Table 4 and Figure 4B). The downregulation of cuticular component is entirely driven by 13 collagens (out of 89 genes in the genome).

**Figure 4.**
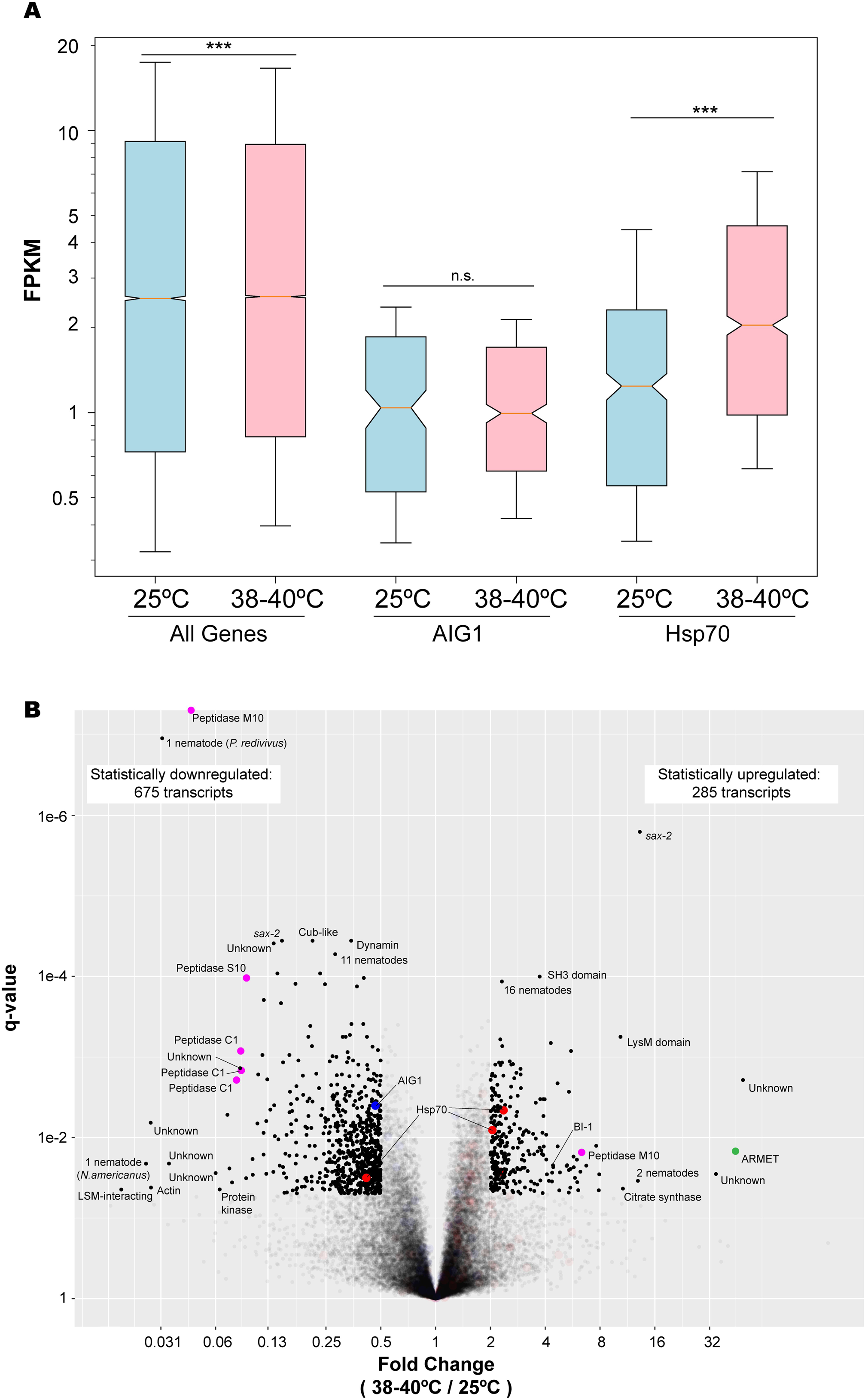
Transcriptome analysis of gene expression in *H. mephisto*. A. Boxplot showing that Hsp70 genes are induced on heat while AIG1 are unchanged. Box shows median and first and third quartiles, while whiskers indicate the 15th to 85th percentiles, and notches represent confidence intervals. P-values obtained by 2-tailed Mann-Whitney test. B. Volcano plot of gene expression fold change under hot (combined replicates: 38°, n=3, and 40°, n=6) versus 25°C control (replicates, n=3). Statistically altered transcripts, defined as q-value less than 0.05 and upregulated or downregulated at least 2-fold under heat-stress conditions (38-40°C) relative to 25°C controls, are indicated with luminosity of 1 while nonsignificant are luminosity 0.1. Genes labeled ‘Unknown’ are novel, as discussed in the text. Proteins encoding no recognizable domain but matching other nematodes by blastp are indicated with the number of nematode species matched (out of the 28 used in constructing the blast database). For genes matching only a single other nematode matches, the species is given in parenthesis. Color key: red: Hsp70, blue: AIG1, green: ARMET, and magenta: peptidases. Abbreviations: Sax-2, Sensory axon guidance 2. LSM, Like SM. BI-1, Bax Inhibitor – 1. ARMET, arginine-rich, mutated in early-stage tumors.

**Table 4.** GO terms enriched in *H. mephisto* genes downregulated under heat stress.

| GO identifier | name | ratio in<br>study | ratio in<br>population | p_uncorrected | p_bonferroni |
| --- | --- | --- | --- | --- | --- |
| GO:0008234 | cysteine-type peptidase activity | 19/649 | 133/28062 | 4.83E-07 | 0.0014 |
| GO:0042302 | structural constituent of cuticle | 13/649 | 89/28062 | 9.91E-07 | 0.0029 |
| GO:0004587 | ornithine-oxo-acid transaminase activity | 3/649 | 3/28062 | 1.23E-05 | 0.0362 |

Hsp70 genes respond to heat (Figure 4A) but with two exceptions they do not make the 2-fold cutoff, and one is actually statistically inhibited on heat stress (Figure 4B). One AIG1 gene is included in the statistically downregulated set (Figure 4B). An intriguing finding is SAX-2 (Sensory AXon guidance 2), a protein involved in neuronal development in *C. elegans* ^56^, and whose inactivation causes a variety of developmental, cellular, and behavioral phenotypes^57^. In *H. mephisto* heat stress causes an isoform switch, with a longer version, MSTRG.2841.1, expressed exclusively at 25°C to a shorter version, MSTRG.2841.2, exclusively expressed at 38-40°C. Thus, *sax-*2 is identified as a transcript both heat-upregulated (12.9-fold) and heat-downregulated (7.0-fold) on the volcano plot (Figure 4B). The long isoform is 9,286 bp and encodes a 3,016 amino acid (aa) protein, with the alternative isoform being 132 basepairs shorter owing to an 84 bp change in transcriptional start site and alternate 3’ splice site choice for exon 10 (of 16 exons total). These isoform differences only result in a 16 aa deletion from the predicted protein. Given that the protein is ∼3,000 aa, and the 16 aa change does not significantly alter the encoded domains (identified by HMMER as MOR2_PAG1 N, mid, and C-terminal domains), the functional implications of the transcriptional shift and alternative splicing of SAX-2 remains to be investigated in future work.

Among the most strongly heat-induced genes was arginine-rich, mutated in early-stage tumors (ARMET), also called mesencephalic astrocyte derived neurotrophic factor (MANF), a gene involved in the unfolded protein response (UPR) in the endoplasmic reticulum (ER)^58^. ARMET was upregulated over 44-fold by heat in *H. mephisto* (Figure 4B). In mouse and human cells, ARMET interacts directly with the ER Hsp70 protein BIP / GRP78^59^. Therefore under hot conditions it may be vital for *H. mephisto* to co-express Hsp70 and ARMET, particularly if they synergize in responding to the damage due to heat in different cellular compartments, Hsp70 in cytosol and ARMET in ER. It appears that *H. mephisto* has developed different genomic strategies for upregulating these two genes: mild upregulation of many paralogous genes (Hsp70) versus extremely highly induced expression of a single-copy gene (ARMET). Regardless, ARMET is a strong marker of ER stress with cytoprotective roles^58,60,61^.

Another upregulated gene was Bax Inhibitor-1 (BI-1), which was upregulated 4-fold (Figure 4B). BI-1 is a conserved anti-apoptotic protein that prevents ER stress-induced apoptosis^62^. BI-1 interacts, binds and suppresses IRE1α activity by cancelling its endoribonuclease and kinase activity activity to promote cell survival^63^. As an ER stress pro-survival factor^62^, BI-1 potentially compensates for the lack of heat induction of AIG1, which plays a similar role inhibiting apoptosis^46^, but may be more specifically tuned to the UPR response.

## Discussion

As the founding genome sequence of the *Halicephalobus* genus, *H. mephisto* is a rich repository of novel biology. *H. mephisto* separated from *Caenorhabditis* at least 22 million years ago^52,64^, and likely over 100 million years ago^65^, though calibrating the nematode molecular clock is difficult because of their poor fossil record^64^. Our data suggest that nematodes access the deep subsurface from surface waters facilitated by seismic activity^66^. This transition from surface to deep subsurface would be expected to exert strong selective pressures on their genomes, which in nematodes are particularly evolutionarily dynamic^6,7, 10–12^. Therefore we speculate that the *H. mephisto* divergence may reflect selection more than neutral genetic drift, making it a particularly informative genome. Consistent with this, the expanded Hsp70 and AIG1 gene families are extremely divergent from earlier exemplars, residing on extended branches in phylogenetic analysis (Figures 2 and 3), and positive selection was detected along several lineages of the Hsp70 phylogeny (Figure 2B).

We investigated the novelty of *H. mephisto*’s 16,186 protein-coding genes by a combination of domain search (HMMER against the Pfam-A database), blastp against the manually curated high-confidence Uniprot-Swisprot database, blastp against the combined proteomes of 28 nematode species (listed in Methods), and Interproscan 5 analysis. At an e-value of 1e-4 we found 10,567 proteins identified by one or more of these methods, leaving 5,619 unknown genes (34.7% of 16,186) lacking domains or any recognizable homology. Nevertheless, 3,599 (64.0%) of the unknown genes are expressed (defined as having at least 5 FPKM across 12 replicates) increasing our confidence they are real genes. Genes whose expression was not detected in our analysis may be expressed at extremely low levels (below 5 FPKM across all replicates) or be expressed under different environmental conditions than the laboratory culture we employed. We conclude that these 5,619 completely unknown genes, and particularly their 3,599 expressed subset, are intriguing candidates for functional adaptation to the deep terrestrial subsurface.

These data are consistent with a previous report that new nematode genomes tend to yield around 33% proteins that are unrecognizable outside their genus^6^, given that *H. mephisto* is the first of its genus to be sequenced fully. As a control we examined the proteome of *Panagrellus redivivus*, using the identical protein-identification pipeline, finding 33.8% unknown genes, quite similar to the numbers for *H. mephisto.* We also tested the pine wood nematode, *Bursaphelenchus xylophilus*, and identified 25.6% unknown genes. Like *H. mephisto, P. redivivus* and *B. xylophilus* are the first genomes of their respective genera to be sequenced. When a within-genus comparison is available, the number of novel genes has been reported to drop to around 10%^6^, which we tested by examining the proteins of *Meloidogyne hapla*, which has a within-genus match to *M. incognita* in the 28 nematode blast database. Consistent with predictions, we identified 8.4% unknown proteins in *M. hapla*. Therefore we conclude that while the genomic plasticity we observe for *H. mephisto* is significant, the number of unknown genes is broadly consonant with nematode molecular systematics showing that roundworm genomes are extremely dynamic. The novelty of the *H. mephisto* genes reflect a combination of evolutionary adaptation and a lack of closely related comparative *Halicephalobus* species in databases. Supporting this, of the 1,730 genes that match nematode genome(s) only, and were not identified by Interproscan, Uniprot-Swissprot, or Pfam, 1,480 of them match *P. redivivius* (Figure S2A), the nearest sequenced nematode relative of *H. mephisto* (Figure 1A). These 1,730 genes are not widely conserved even among roundworms, with only 17 (1%) of them identified in all 28 species and most (511) identified in only one other species (Figure S2B); unsurprisingly the majority of these (399, 78%) were only found in *P. redivivus*.

The assembled genome of *H. mephisto* is smaller (61.4 Mb) than most other sequenced nematode genomes with the exception of *M. hapla* (54 Mb)^67^ and *M. incognita* (47-51 Mb)^68^, though similar in size to the most closely related species, *P. redivivus* (64.4 Mb)^18^. *H. mephisto* reproduces via parthenogenesis^1^, while both *M. hapla* and *M. incognita* are facultatively parthenogenetic^67,68^. In *Caenorhabditis,* loss of males has been linked to genome shrinkage^11^. In *M. incognita*^68^ and *D. pachys*^69^ the loss of sexual reproduction leads to functional haploidization as alleles diverge into paralogs across the genome (leading to most genes being present as duplicate, divergent copies). The conversion of a diploid into a functional haploid genome is associated with three predominant changes: 1) a high degree of heterozygosity as lack of recombination leads to high divergence between alleles, now paralogs, 2) assembly of a haploid genome, and 3) detection of two copies of most genes that are single-copy in other organisms^68,69^.

The heterozygosity we observed in *H. mephisto* is modest: The kmer frequency distribution from Illumina reads shows two prominent peaks consistent with approximately 1% heterozygosity^70^ (Figure S3A) and mapping reads back to the assembly identified 707,190 snps and 55,683 indels (762,873 total variants) confirming an overall snp heterozygosity of 1.15%. In contrast, *D. pachys* displays 4% heterozygosity^69^.

The *H. mephisto* genome assembly we obtained is largely haplotype-merged: mapping the reads back to the genome shows that 59.7 Mb of the assembled sequence (97%) is at haplotype-merged coverage (102x) and only 1.7 Mb (3%) exists as potentially diverged haplotypes at lower coverage (Figure S3B). In agreement, CEGMA analysis of assembled reads (Figure S3C) and fragments (Figure S3D) are predominantly single peaks at approximately 100x coverage.

Allele-to-paralog conversion results in two recognizably distinct gene copies, or paralogs, in the genome. In the *D. pachys* genome CEGMA found an average of 2.12 copies of each core eukaryotic ortholog^69^ yet in *H. mephisto* CEGMA^13,14^ reported an average 1.19 copies of each gene while BUSCO^16^ identified 97% single-copy genes. We therefore conclude that *H. mephisto* exhibits very little functional haploidization into paralogs, and it may represent an early evolutionary stage in this process.

It is important to verify that amplified Hsp70 and AIG1 gene families do not represent assembly errors. One way this could happen is if an assembler produces multiple overlapping contigs encoding the same locus, leading to artifactually high gene copy numbers. In that case multiple near-identical copies of the genes would be present. To test this we extracted the corresponding nucleotide sequences for each family and performed within-family, all-vs-all blastn at an evalue of 1e-4 and filtered for non-self matches. These nonself matches were relatively divergent: the best nonself within-family blastn match for 112 Hsp70 loci averaged 86.4% identity at the nucleotide level (Figure S4A) and for 63 AIG1 loci averaged 89.7% identity (Figure S4B). Given the observed genomic heterozygosity is only 1.15%, these data suggest that the Hsp70 and AIG1 genes are diverged paralogs and neither allelic copies nor redundant misassemblies. As a control we extracted 65 collagen genes and performed the same analysis. As might be predicted for an ancient and highly divergent gene family, 74% of collagen genes lacked within-family nonself blast matches entirely, though those that matched averaged 86.6%, similar to Hsp70 (Figure S4C). These sequence divergences are greater than a control analysis performed by blast of the assembly to itself (Figure S4D). We conclude that the expanded Hsp70 and AIG1 families represent parology rather than assembly redundancy artifact.

Often repetitive elements are collapsed by assemblers, so we checked that Hsp70 and AIG1 are not collapsed repeats, which would cause under-representation of their family diversity. Mapping raw reads onto the assembled genome indicates the collapsed repeats as regions of elevated coverage^71^. By mapping the raw reads back to the *H. mephisto* genome we found that the coverage of both Hsp70 and AIG1 families are not elevated relative to the entire genome (Figure S3E). Overall we conclude that the Hsp70 and AIG1 genes identified in this study are true paralogs, neither over nor under-represented in the genome.

*D. pachys* is remarkable for having fused its chromosomes together into a single linkage group and eliminating telomeres. However, over 6,000 telomeric repeat-containing reads (at least 4 copies of TTAGGC, the *C. elegans* telomeric repeat^72^) were present in the raw Illumina data from *H. mephisto.* By extracting read pairs with telomeric repeats in at least one read, merging them with PEAR^73^, and assembling them with MIRA we were able to identify 7 unique subtelomeric regions, suggesting that while the number of *H. mephisto* chromosomes may be reduced relative to *C. elegans*, they are not fused as in *D. pachys*. Consistent with this, we find two homologs of the telomeric gene Protection of Telomeres (POT1) in *H. mephisto* relative to the three in *C. elegans,* all of which are lost in *D. pachys*^69^. Also in contrast to *D. pachys,* we were able to identify Telomerase Reverse Transcriptase (TERT) in *H. mephisto*, suggesting that standard telomeres have been retained in the subterrestrial organism and chromosome fusion is not an inevitable consequence of a parthenogenetic lifestyle.

Gene expression provides clues to the adaptation to the warm subterrestrial environment. Among genes whose expression change significantly on exposure to heat, over twice as many are downregulated (675) as upregulated (285) (Figure 4B). We suggest two potential explanations of this phenomenon: the worms may find lower temperatures more stressful given their native conditions are warmer, and therefore activate more genes at 25°C relative to 38-40°C. Or, the downregulation of genes may be itself an adaptation to heat, an idea consistent with the Regulated IRE1α Dependent Decay (RIDD) pathway of the Unfolded Protein Response (UPR)^74,75^. When the RIDD pathway is activated, degradation of ribosome-bound transcripts is mediated by the endoribonuclease domain of IRE1α^74^, relieving the immediate protein synthesis demand on the endoplasmic reticulum, and providing existing proteins time to re-fold^63,74^. Our data cannot definitively distinguish these two theories; however, the elevated expression of protein chaperones like Hsp70 under heat exposure (Figure 4A) supports the model in which observed changes in transcriptional profile reflect an adaptive response to higher, rather than lower, temperatures. Combined with the observed heat-induced expression of the anti-apoptotic factor Bax Inhibitor 1 (BI-1) (Figure 4B) this response would help *H. mephisto* survive the abiotic heat stress of the subterrestrial environment.

While we report significantly expanded Hsp70 and AIG1 families, only the Hsp70 genes are upregulated under heat stress in the laboratory (Figure 4A). Interestingly the worms appear to keep per-gene Hsp70 expression low, even under conditions of heat stress. The per-gene expression of Hsp70 at high temperature is slightly lower than all genes (median Hsp70 FPKM = 2.04, and 2.58 for all genes, Figure 4A), but the dramatic expansion of Hsp70 paralogs effectively elevates the gene dosage for Hsp70 to 51-fold higher than a single-copy gene would be. This is a similar order of induction as the ARMET protein, an UPR-related single-copy gene induced 44-fold (Figure 4B), and which may synergize with Hsp70. While many studies have shown induction of Hsp70 genes upon heat-shock, the short-term exposure of non-heat-adapted organism to brief extreme heat^31–34^, organisms under long-term adaptation to heat tend to minimize overexpression of Hsp70 because it has harmful effects on development, fertility, and growth^29,76–78^. Long-term exposure to heat stress led to downregulated Hsp70 expression in both flies^79^ and fish^80^. The subterrestrial environment of *H. mephisto* is thermostable over time: four readings during a five-year period showed the water temperature was an average of 36.8 +/- 1.2°C^2–4,81,82^. Similarly, we cultured the worms for 2-4 weeks at constant temperatures (25°C, 38°C, or 40°C) in the laboratory for RNA isolation and gene expression analysis. Therefore, *H. mephisto*’s sustained expression of Hsp70 under conditions of constant stable heat stress implies a functional bypass of the genes’ known detrimental effects on growth and development^29^. We hypothesize that the divergent Hsp70 genes in *H. mephisto* that respond most strongly to elevated temperatures may have been functionally modified to ameliorate these deleterious effects, marking them as important candidates for future study.

We were surprised to find that the expansion of AIG1 in *H. mephisto* likely mediates abiotic stress aside from heat. These genes respond to abiotic stress, including heat, in Arabidopsis^48^ and in mammals the proteins are involved in immune system function, inhibiting apoptosis during T-cell maturation^45^. Given that *H. mephisto* did not activate AIG1 under heat stress, we posit these genes are involved in responding to hypoxia or other abiotic non-thermal stresses present in the deep terrestrial subsurface. Remarkably, however, heat induced another anti-apoptotic factor in *H. mephisto*--the Bax Inhibitor 1, BI-1^62^, suggesting a different method for blocking apoptotic response under subterrestrial heat stress. Together with our work, these data suggest *H. mephisto* has adapted to the subterrestrial environment by managing unfolded protein stresses while upregulating Hsp70 and inhibiting apoptosis.

We note that the pacific oyster *Crassostrea gigas* has convergently expanded Hsp70 and AIG1 gene families^83^ and activates the UPR in response to abiotic stress including heat^84^, so *H. mephisto* helps define a general evolutionary adaptive response to heat stress. While the oyster experiences considerable thermal fluctuation, *H. mephisto* does not, as described above. Therefore, the signature of adaptation we report is not limited to cyclical or temporary temperature fluctuations but extends to adaptation to constant warm environments.

The expansion of Hsp70 is shared, also convergently, by distantly related *Diploscapter* species, soil nematodes which display pronounced thermotolerance^37,39^, and we show here that positive selection has driven Hsp70 expansion in *Diploscapter* and *H. mephisto*.These findings may not only relate to environmental heat: *D. coronatus* has been reported as a facultative parasite of humans, so it must survive to 37°C, human body temperature^38^. Consistent with this, the closest relative of *H. mephisto* is a deadly horse parasite, *H. gingivalis*, which has not been fully sequenced, and has also been reported as a facultative and fatal parasite of humans^85^. Therefore, the genome signature of adapation to heat in *H. mephisto*, *D. coronatus,* and *D. pachys* may serve as a preadaptational bridge to parasitic lifestyles at least in some lineages. Therefore these genomic adaptive strategies are of significant concern to human and animal health, and as our climate warms, it will be increasingly important to understand their evolutionary dynamics.

## Acknowledgements

The authors acknowledge Laura Landweber, in whose laboratory J.R.B prepared the *H. mephisto* Illumina DNA library. We thank Wei Wang, Jessica Wiggins, and Donna Storton of the Princeton Sequencing Core Facility for assistance with Illumina library preparation and sequencing. We also acknowledge Aaron Goldman for early work done on the *H. mephisto* project, and Elizabeth Ginsburg for comments on the manuscript. We thank Evgeny Bisk for assistance with the installation and troubleshooting of software on the Zorro Computer Cluster at American University. Mariana Erasmus acknowledges support from the Technology Innovation Agency (TIA), an agency of the Department of Science and Technology (DST) and is grateful to Izelle de Beer and Bernice Jordaan for their assistance in culturing of the nematodes.We are indebted to the logistical support of Sibanye Gold Limited, the management and staff of Beatrix Gold Mine that enabled the capture of *H. mephisto* and follow up samples for metagenomic studies. The work of M.C.Y.L was supported by NSF Grant #DEB-1441646 to T.C.O, and J.R.B was supported by NIH 1K22CA184297 and a NASA DC Space Grant Consortium grant which funded the RNA and PacBio sequencing.

## Author Contributions

D.J.W., S.E.A. and J.R.B. performed gene prediction and phylgenetic tree construction; D.J.W. and J.R.B. performed transcriptome analysis and wrote the paper, and J.RB. performed genome assembly. M.C.Y.L constructed the multi-locus phylogenetic tree. M.E. and E.v.H. grew the nematodes for RNA. K.C.A. performed analysis of positive selection and K.W.C performed repetitive element analysis. G.D. and R.S performed Illumina RNA and PacBio DNA sequencing. G.B. cultured the nematodes for DNA extractions and with T.C.O. provided critical feedback on the manuscript drafts and input into experimental design.

---

**Figure S1.**
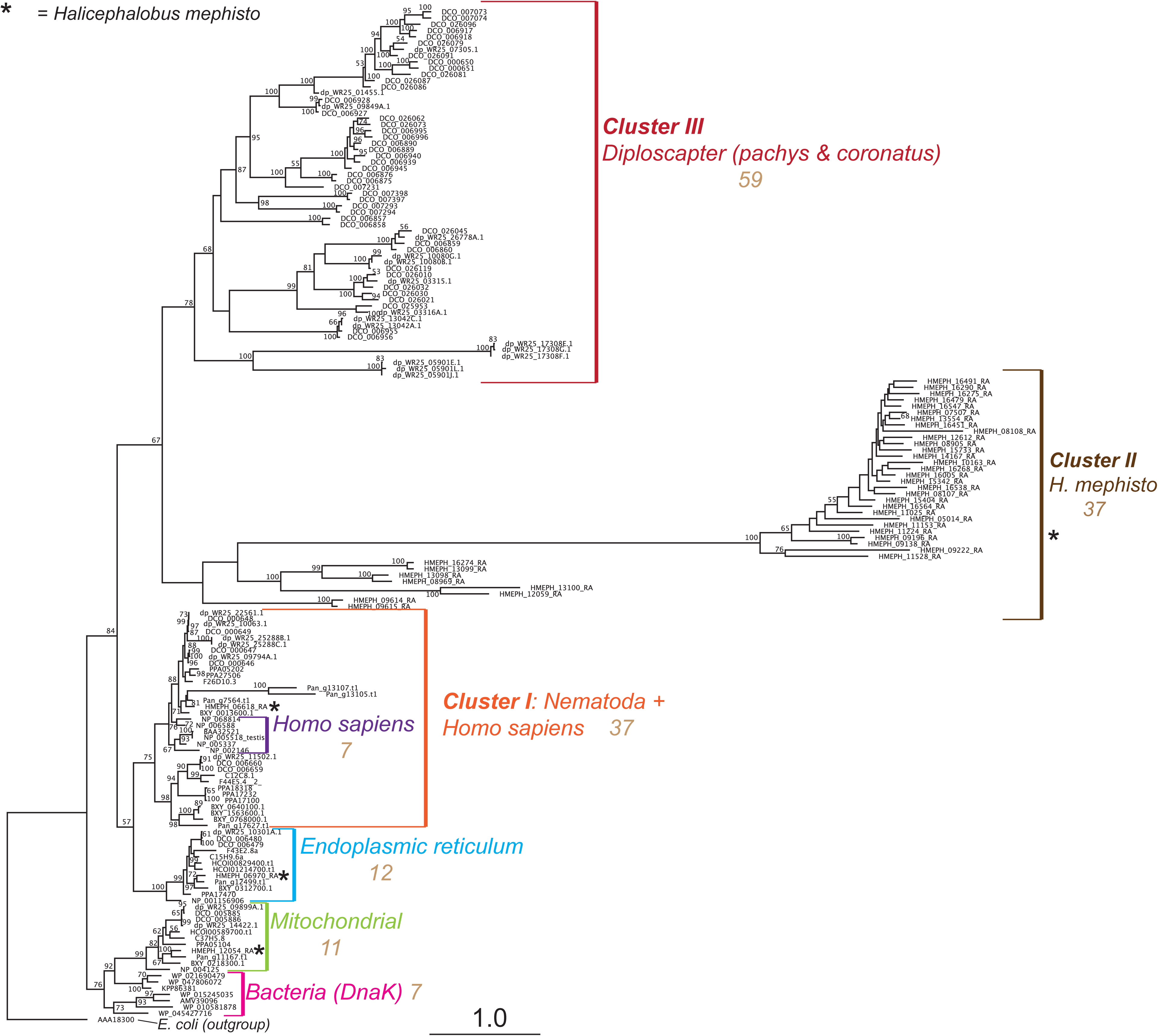
RAxML phylgenetic tree of Hsp70 domains. *H. mephisto* sequences marked with an asterisk (*) and *D. pachys* with arrows. Branch numbers indicate bootstrap support, scale bar represents substitutions per site.

**Figure S2.**
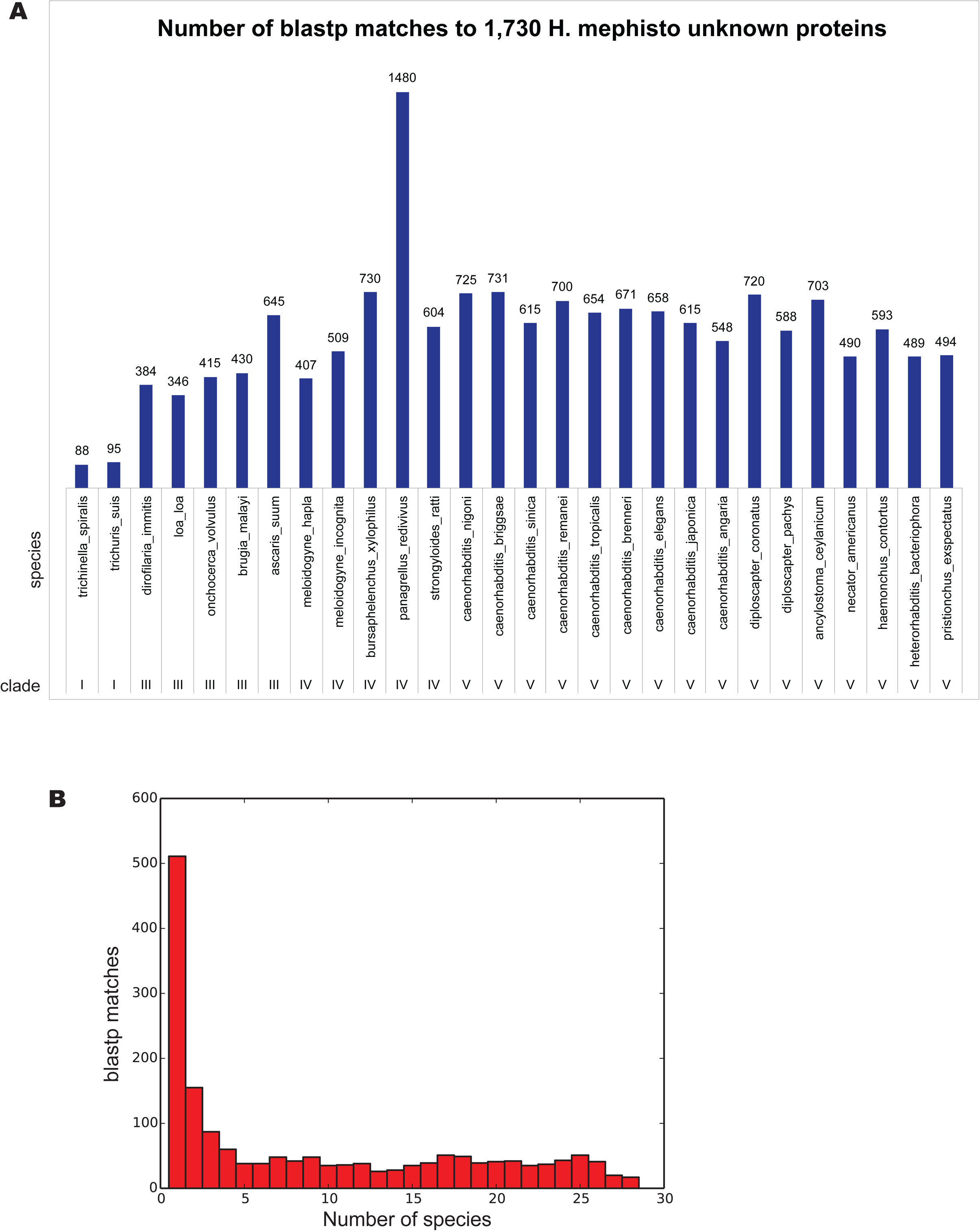
A. Analysis of 1,730 *H. mephisto* genes matching to at least one nematode proteome but otherwise lacking recognizable domains by Interproscan or Pfam, and lacking matches to Uniprot-Swissprot. The evalue threshold was set to 1e-4 and number of matches per species are plotted. B. Histogram showing the number of species matched for each of the 1,730 intra-nematode proteins.

**Figure S3.**
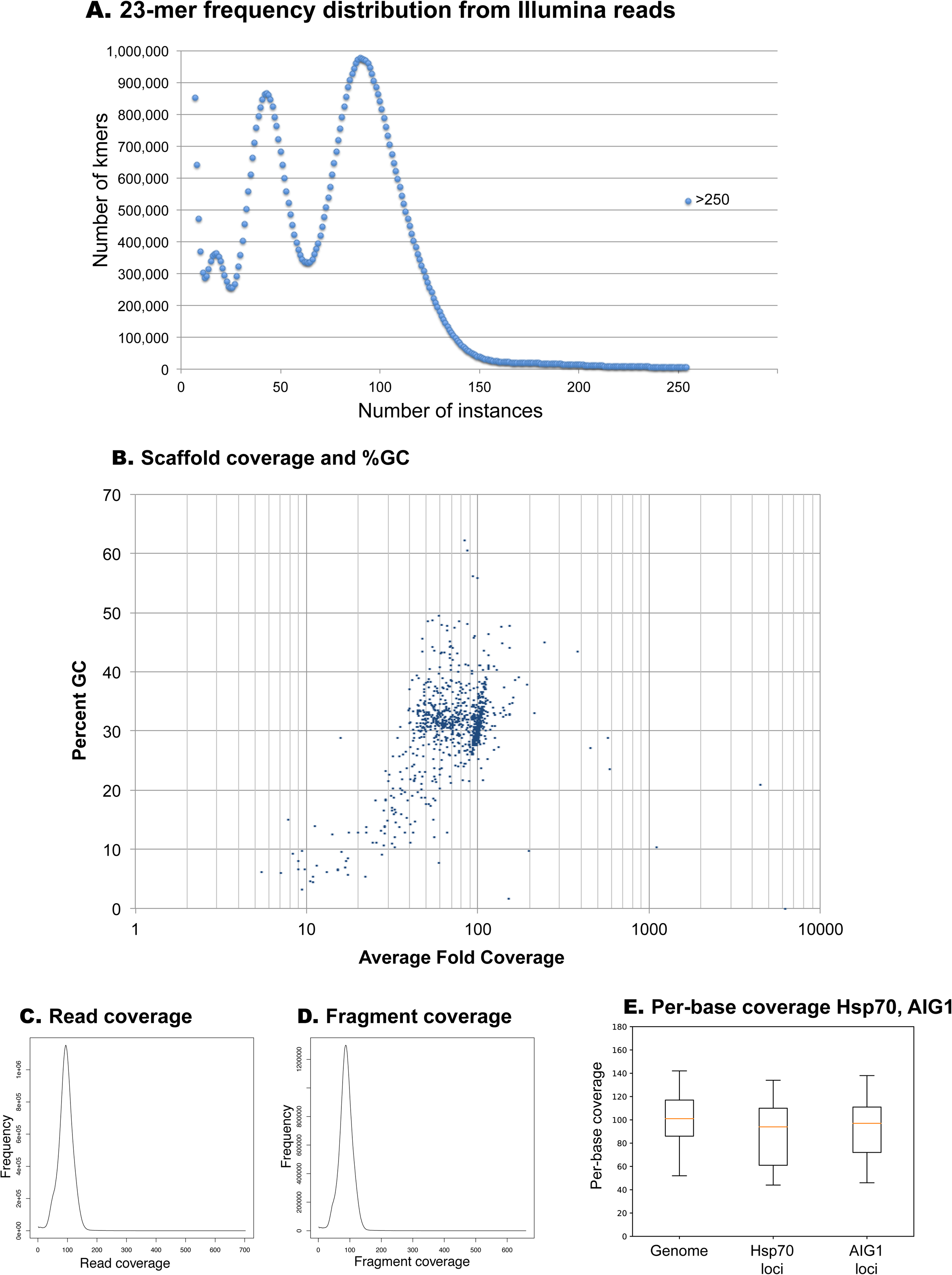
K-mer frequency and assembly coverage. A. Kmer frequency distribution (23-mers). B. Per scaffold coverage vs percent GC. C. Assembled read coverage from CEGMA. D. Assembled fragment coverage from CEGMA. E. Boxplot of per-base coverage of the *H. mephisto* genome, Hsp70 loci, and AIG1 loci.

**Figure S4.**
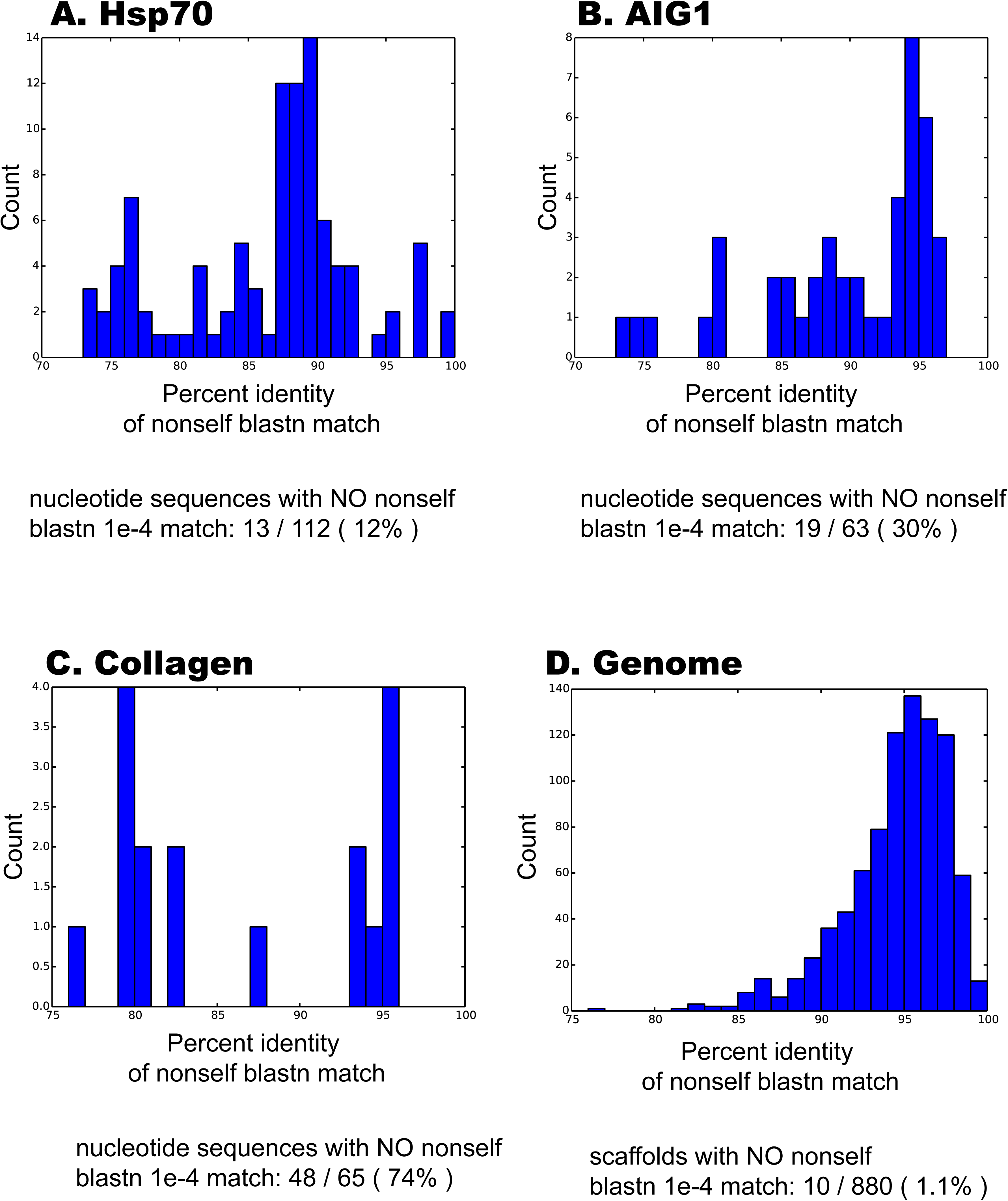
Analysis of degree of similarity at nucleotide level between members of expanded gene families. Self-self blastn was performed with evalue 1e-4, then the percent identity of the first non-self match was plotted. A. Hsp70, B, AIG1, C, Collagen (control), D, full genome (control).

## Methods

### *H. mephisto* culture and isolation of DNA and RNA

*H. mephisto* were cultured by standard *C. elegans* methods^86^, on agar plates seeded with *Escherichia coli* OP50. For DNA extraction the NucleoSpin kit (Cat #740952.250, Macherey-Nagel, Bethlehem, PA, USA) was used. The pellet was resuspended in 540 µl T1 buffer supplemented with 10 µl Proteinase K. Lysis was accomplished by 4 cycles of rapid freeze-thaw using a dry ice-ethanol bath, with thawing on a 56°C heatblock; cycles were approximately 1 minute per freeze or thaw step. After this the sample was treated with an additional 25 µl proteinase K overnight at 56°C. After this the manufacturer’s protocol was followed for column purification of high-quality DNA, which was verified by gel electrophoresis prior to library construction.

For RNA, the worms were cultured on 5% agar plates at 25°C, 38°C, or 40°C for 2-4 weeks prior to harvest, then pelleted in PBS pH 7.7, and flash-frozen or immersed in DNA/RNA Shield Buffer for storage and extraction of nucleic acids. For RNA extraction Zymo’s Duet DNA/RNA MiniPrep Plus kit (Cat # D7003) was used. The worm pellet was resuspended in 300µl of DNA/RNA Shield buffer, transferred to a tube of 0.5mm BashingBeads (Zymo Cat # S6002), and homogenized on a vortexer at maximum speed twice for 5 minutes with a 1-2 minute rest period between (for cooling). After this 30 µl of PK buffer and 15 µl Proteinase K were added, and the solution incubated for 30 minutes at 55°C, then supplemented with 345 µl lysis buffer. After pelleting the insoluble material at 16,000 g for 1 min, the supernatant was transferred to yellow (for DNA) and green (for RNA) columns as described in manufacturer’s protocol for the Duet DNA/RNA MiniPrep Plus Kit, performing in-column DNAse treatment of RNA as recommended.

Based on Agilent BioAnalyzer 2100 output, a total of 12 samples yielded RNA of sufficient quality for sequencing: 3 from 25°C, 3 from 38°C, and 6 from 40°C. For analysis the 38°C and 40°C samples were considered together as ‘high’ temperature and the 25°C replicates as ‘normal’ temperature.

### Genomic DNA Sequencing

For Illumina, a TruSeq library with insert size of 387 bp was generated with 9 cycles of PCR after gel purification. This library was paired-end sequenced on a HiSeq2500 with 215 bp reads, yielding 58.7 million pairs. This data was assembled with Platanus^70^ as described in the next section. For PacBio, 30 lanes were run on the RS II system using libraries generated off the same DNA sample used in Illumina.

### RNA Sequencing

Stranded RNA-seq libraries were generated using the KAPA Total Stranded preparation kit (KAPA Biosystems catalog # KK8484) with an average insert size of 175 bp. Samples were sequenced on a HiSeq 2500 in High Output mode using 2×50 bp paired-end reads, generating at least 60M total reads per sample. Preliminary quality control for read mapping was performed using taxMaps version 0.2.1^87^, on a downsampled subset (1%) with the NCBI BLAST nt and a kmer size of 75 to confirm species etiology for the reads generated.

### Genome Assembly

Raw read kmer analysis was performed with the SoapEC v. 2.01 (from the SOAPdenovo2 package^88^), KmerFreq_HA set to kmer 23 and error corrected with Corrector_HA. We found that platanus assembly was optimial at 100x average coverage, so a total of 15,582,039 random paired, error-corrected reads were used in the final assembly. Platanus version 1.2.4^70^ was used to assemble with a stepsize 2, kmer of 21, and -u 1. Platanus scaffolding was performed with settings -n 345, -a 386, and -d 62. Subsequently gaps were closed with Platanus gapclose. Scaffolds less than 500 bp were discarded and this assembly was further scaffolded with 30 lanes of PacBio data using the PBJelly component of PBSuite v. 15.8.24. Reapr v. 1.0.18 (perfectmap, -b) was used to break chimeras and erroneous gaps. We removed sequences under 1000 bp and identified 40 bacterial scaffolds by a combination of coverage (less than 26X) and GC content (over 55%), which blast confirmed as prokaryotic. These sequences were found to encode a complete sphingomonas genome to be described elsewhere, but does not appear to be a deep subterrestrial inhabitant based on metagenomic borehole read mapping. After additional removal of the mitochondrial scaffold, this final *H. mephisto* nuclear genome of 880 scaffolds was used in all further analysis. This assembly has an N50 of 313kb, with the longest 2.55 Mb and is highly contiguous: the final assembly encodes only 10 gaps encoding 476 bp (0.0008% of sequence).

### Heterozygosity

The error corrected reads were mapped back to the final *H. mephisto* assembly with bwa-mem v.0.7.12 and mis-mapped reads and PCR duplicates were removed with samtools v. 1.9. Using the remaining 31,867,988 reads, snp variants were called with bcftools v. 1.9 mpileup and then call command with the -mv flag.

### Analysis of within-family non-self blastn matches

Gene family coding sequences were extracted into a fasta file based on the transcript coordinates from the GFF file produced by gene annotation with Maker2 and Stringtie. These fasta files were used to build blast databases. These databases were each queried using blastn (at 1e-4) using the same fasta file as query that was used to build the database (thus, performing all-vs-all blastn). The resultant blast output in tabular format (-outfmt 6) was parsed using a custom python script to isolate only the first (best) non-self match from the blast output, and a histogram was generated of the percent identies of these non-self matches. For the genomic assembly all-vs-all comparison, the same process was carried out but using the 880-contig full genome assembly.

### Analysis of coverage

Genome-wide coverage was calculated using the samtools v. 1.9 depth command on the bamfile generated for heterozygosity analysis, followed by custom parsing of the coverage file with a python script. The same coverage file was parsed using the unique transcript coordinates as described in the “Analysis of within-family non-self blastn matches” section. The per-basepair coverage values for Hsp70, AIG1, and the entire genome were evaluated with custom python script producing a boxplot shown in the figure.

### Analysis of Repetitive Sequence

RepeatModeler^89^ v.1.0.11 was used to create a custom repeat library. This library was screened for accuracy with HMMER^90^ v 3.1b2 to identify mis-classifed protein-coding genes, which were removed. This library was used in a RepeatMasker^89^ v 4.0.6 run using the default parameters. The initial RepeatMasker run designated 21.07% of the genome as consisting of transposable elements, of which 18.48% of the total genome, or 87.7% of the repeat segments, as unclassified repeats. Subsequently, nhmmer^19^ analysis was run on the identified repeats using the DFAM database^91^ with e-value set to 1e^−2^ to accommodate the highly divergent genome.

### Gene discovery

Maker2^20^ version 2.38.1 was utilized to run Augustus and SNAP as *ab initio* predictors to make comprehensive gene predictions for *H. mephisto,* and incorporating 28 nematode proteomes as hints along with the RNA-seq data. These gene predictions were refined utilizing Tophat2-StringTie-Ballgown suites of programs^21,22^, which also estimate expression levels. Tophat2 v.2.1.1 was used to align the RNA-seq data against the *H. mephisto* genome with Maker2 predicted genes as a input .gff3 file. The resulting .bam files were fed into StringTie v. 1.3.4, to generate a transcriptome annotation of each, as well as quantify the expression levels and estimate the abundance of each transcript, which were subsequently unified using Stringtie’s merge function^22^. Ballgown v. 2.12.0 plots the gene abundance and expression data for visualization, from the StringTie output data^22^.

Together StringTie and Maker2 predicted 34,605 transcripts across 12 different RNAseq datasets, which map to a distinct set of 17,209 unique loci as defined by gffcompare. From these loci the longest protein sequence predicted by TransDecoder^23^ v. 5.3.0, was used in domain comparisons with *C. elegans,* the reference proteome UP000001940_6239 (19,922 nonredundant proteins) from Ensembl RELEASE 2018_04. TransDecoder was run in strict mode, requiring at least 50 amino acids, and only the single best cds prediction per transcript retained, to obtain the nonredundant set of 16,186 protein-coding genes.

The 28 nematode proteomes used are as follows (all obtained from WormBase Parasite, https://parasite.wormbase.org):

ancylostoma_ceylanicum.PRJNA231479.WBPS12.protein.fa
ascaris_suum.PRJNA62057.WBPS12.protein.fa
brugia_malayi.PRJNA10729.WBPS12.protein.fa
bursaphelenchus_xylophilus.PRJEA64437.WBPS12.protein.fa
caenorhabditis_angaria.PRJNA51225.WBPS12.protein.fa
caenorhabditis_brenneri.PRJNA20035.WBPS12.protein.fa
caenorhabditis_briggsae.PRJNA10731.WBPS12.protein.fa
caenorhabditis_elegans.PRJNA13758.WBPS12.protein.fa
caenorhabditis_japonica.PRJNA12591.WBPS12.protein.fa
caenorhabditis_nigoni.PRJNA384657.WBPS12.protein.fa
caenorhabditis_remanei.PRJNA248909.WBPS12.protein.fa
caenorhabditis_sinica.PRJNA194557.WBPS12.protein.fa
caenorhabditis_tropicalis.PRJNA53597.WBPS12.protein.fa
diploscapter_coronatus.PRJDB3143.WBPS12.protein.fa
diploscapter_pachys.PRJNA280107.WBPS12.protein.fa
dirofilaria_immitis.PRJEB1797.WBPS12.protein.fa
haemonchus_contortus.PRJEB506.WBPS12.protein.fa
heterorhabditis_bacteriophora.PRJNA13977.WBPS12.protein.fa
loa_loa.PRJNA246086.WBPS12.protein.fa
meloidogyne_hapla.PRJNA29083.WBPS12.protein.fa
meloidogyne_incognita.PRJEB8714.WBPS12.protein.fa
necator_americanus.PRJNA72135.WBPS12.protein.fa
onchocerca_volvulus.PRJEB513.WBPS12.protein.fa
panagrellus_redivivus.PRJNA186477.WBPS12.protein.fa
pristionchus_exspectatus.PRJEB6009.WBPS12.protein.fa
strongyloides_ratti.PRJEB125.WBPS12.protein.fa
trichinella_spiralis.PRJNA12603.WBPS12.protein.fa
trichuris_suis.PRJNA179528.WBPS12.protein.fa

## Gene Expression

After gene discovery, expression analysis was carried out using Ballgown^92^ v. 2.12.0 following the protocol as described^22^. Genes with less than 5 total reads across all 12 replicates were filtered from the analysis. Replicates were grouped into ‘high’ (38-40°C) or ‘low’ (25°C) and the Ballgown stattest() function used to identify those genes statistically different between high and low temperatures. The output of stattest() include q-value and fold-change and these were used to generate the volcano plot as a scatterplot with ggplot2 in R. Significantly heat-regulated genes were defined as exhibiting a q-value less than 0.05 and upregulated or downregulated at least 2-fold under heat-stress conditions (38-40°C) relative to 25°C controls. For the boxplots of gene expression the FPKM values at high or low temperature were exported as a text file and imported into Python 2.7.14 where a custom script was used to construct the boxplots with matplotlib 2.2.2.

### Analysis of unknown genes

*H. mephisto* genes were analyzed by blastp (evalue 1e-4) against a collection of 28 nematodes as outlined for Gene Discovery. When blasting with controls *P. redivivus, M. hapla,* or *B. xylophilus* we created a separate database removing itself (to avoid the trivial self-matching) and replaced it with the *H. mephisto* proteome, keeping a 28 nematode species comparison set. For all analyses we also performed blastp against the uniprot-swissprot manually curated database (1e-4), Hmmer domain search against the PfamA database (1e-4), and Interproscan 5.30-69.0 running TIGRFAM 15.0, Hamap 2018_03, SMART 7.1, PRINTS 42.0, and Pfam 31.0. Custom python scripts were used to combine output of all analyses and identify true unknown genes.

### Venn Diagram

The genome of each species was uploaded onto the selected template on the website OrthoVenn (http://www.bioinfogenome.net/OrthoVenn/).

### Domain Comparisons

HMMER^93^ was used to identify protein domains from both *H. mephisto* and *C. elegans* nonredundant protein predictions with evalue 1e-10. Domain counts of *H. mephisto* were compared to *C. elegans* using a custom python script.

### Multi-locus phylogenetic tree

Phylogenetic relatedness of *Halicephalobus mephisto* relative to other nematode species in Clade IV was determined by constructing a maximum likelihood tree of single-copy orthologous genes. Protein-coding sequences of 21 nematodes were downloaded from WormBase (release WBPS9) on March 26, 2018. They are *Bursaphelenchus xylophilus, Ditylenchus destructor, Globodera pallida, Globodera rostochiensis, Meloidogyne floridensis, Meloidogyne hapla, Panagrellus redivivus, Parastrongyloides trichosuri, Rhabditophanes sp. KR3021, Steinernema carpocapsae, S. feltiae, S. glaseri, S. monticolum, S. scapterisci, Strongyloides papillosus, S. ratti, S. stercoralis,* and *S. venezuelensis*. The three outgroups were Clade V nematode proteomes, namely *Diploscapter pachys* PF1309 (NCBI project number PRJNA280107), *Heterorhabditis bacteriophora* (NCBI project number PRJNA438576), and *Caenorhabditis elegans* (WormBase release WS264). Following the procedures of OrthoMCL v2.0.9^94,95^, the proteome files were modified to the required format by OrthoMCL (step5), and then filtered to remove sequences that are shorter than 10 amino acid residues and have less than 20% of stop codons (step6). An All-verse-All BLAST^96^ search amongst all proteomes was performed as suggested by OrthoMCL step7, which involved creating a BLAST-searchable protein database with masking information, followed by a BLASTp search with an e-value threshold of 1e-5 (“-evalue 1e-5”), and results were stored in a tab-delimited file (“-outfmt 6”). The BLASTp results were parsed using orthAgogue v1.0.3^97^, which filtered out protein pairs with overlap less than 50% (-o 50) and BLAST (or bit) scores below 50 (-u) before identifying valid protein pairs. the single-copy orthologous groups shared by the 22 proteomes. This step substituted OrthoMCL step8 to step11. The resultant orthologs.abc file was then used for clustering (inflation index, -I = 2.0) and creating orthologous groups (OrthoMCL step 12-13). Orthologous groups that contained a single protein sequence from each of the 22 genomes were considered as single-copy orthologous groups (SCOGs). Sequences of 99 identified SCOGs were aligned individually using MUSCLE v3.8.31^98^ and the default parameters, and trimmed using trimAl^99^ to remove residual positions that were shared by less than 50% of the sequences in the multiple sequence alignments. Trimmed alignments were manually examined to make sure there was no spurious sequences or poorly aligned regions, and each was evaluated by ProtTest v3.4.2^100^ to identify for the best substitution model. Sequences of these 99 SCOGs were concatenated by taxa. Using this final multiple sequence alignment of 43,188 amino acid positions (including 34,549 distinct patterns) and the specific best substitution model identified for individual SCOGs, partitioned phylogenetic analysis was performed using raxml-ng v0.4.1b to find the best maximum-likelihood tree. The substitution models used were JTT+G, JTT+G+F, JTT+I+G, LG+G, LG+G+F, LG+I+G, LG+I+G+F, RtREV+I+G+F, VT+I+G, WAG+I+G, and WAG+I+G+F. Robustness of tree topology was evaluated by 100 iterations of bootstrap analysis.

### Go Analysis

The find_enrichment.py script from the GOATOOLS v0.6.10 package^101^ was used under default settings to examine the 285 upregulated and 675 downregulated genes relative to the entire set of proteins. GO terms were assigned using Interproscan 5.30.69 ^102^.

### Hsp70 and AIG1 Tree Building

For Hsp70 full-length proteins were aligned, and for AIG1 the proteins were broken into domains using the envelope coordinates provided by HMMER and a custom Python script. We labeled the domains by order within the original protein. Alignments performed with MAFFT^103^ v.7.017 and refined manually to minimize indels. Trees were generated with MrBayes^104^ v.3.2.6, using the blosum rate matrix and invgamma rate variation; and RAxML 8.2.12^105^ with the PROTCATBLOSUM62 rate matrix and 200 bootstrap replicates. All non-*mephisto* sequences in Figure 2 are identified using their Wormbase ParaSite (https://parasite.wormbase.org/index.html) gene identifier. (For a list of species downloaded from Wormbase, see Gene Discovery section of Methods). For Figure 3 we prepended non-*mephisto* sequences with the shortest possible (normally two letter) genus and species abbreviation (*e.g.,* ‘Dp’ for *Diploscapter pachys*) followed by ‘^’ prior to the Wormbase ParaSite gene identitifer. Non-nematode gene sequences are indicated by their NCBI accession numbers and UniProtKB identifier for *Rhizophagus irregularis*.

### PAML with branch-sites model

Using Codeml (PAML v.4.9)^41,106^ branch site ω was estimated using runmode = 0, seqtype = 1, model = 2, NSsites = 2, in the Codeml control file. Each branch was estimated twice: once with a neutral model (fix_omega = 1 and omega = 1) and once using a purifying selection model (fix_omega = 0, omega = 1). The p values were determined using the likelihood ratio test (LRT) statistic *2Δl* compared against χ^2^ with critical values of 3.84, 5% significance level, and 6.63, for 1% significance^107^.

### Analysis of subtelomeric sequences

Extraction of telomere repeat-containing read pairs (at least one of which contains at least 4 copies of TTAGGC) from raw Illumina data was performed using a custom Python script. These read pairs were merged at their overlaps using PEAR^73^ v0.9.8, and the resultant fused reads were assembled with MIRA^108^ v4.0. The resulting contigs were manually inspected and redundant sequences were collapsed to obtain an estimate of subtelomeric region number.

## Data Availability

This Whole Genome Shotgun project has been deposited at DDBJ/ENA/GenBank under the accession SWDT00000000. The version described in this paper is version SWDT01000000.

The raw Illumina DNA and RNA data, and PacBio DNA data are available on the Sequence Reads Archive (SRA) at accession PRJNA528747.

Transcriptome expression data as FPKM values genome-wide is available in the Gene Expression Omnibus (GEO) at accession GSE133178.

The genome annotation file and transcript and protein predictions are available from WormBase release 14 (WBPS14) onwards as accession Halicephalobus_mephisto_prjna528747.

